# A Characterization of the Immune Cells in Immunocompetent and Immunodeficient Mice with Orthotopic Brain Tumors

**DOI:** 10.1101/2025.04.29.651335

**Authors:** B Gardam, E Nam, PM Kollis, J Kilyen-Coles, S Lenin, BL Gliddon, MN Tea, SM Pitson, MP Brown, LM Ebert, T Gargett

## Abstract

**Background:** Glioblastoma is characterized by poor survival with few treatment advances for over 20 years. Active research is being conducted in immunotherapies; however, this may be hampered by the lack of detailed characterization of the immune compartments of orthotopic murine brain tumor models.

**Methods:** We used cell lines derived from glioblastoma patients and murine glioma cell lines to produce intracranial xenograft and syngraft brain tumors in NSG or C57BL/6 mice, respectively. High-parameter flow cytometry and tissue immunofluorescence staining were used to characterize the immune cell compartment in each model. We investigated brain, spleen, lymph node, and bone marrow for the frequencies of different immune cells. Further, we investigated the effect of sub-lethal whole-body irradiation, commonly used for lymphodepletion, on immune cells in immunocompetent mice.

**Results:** In our study of non-tumor-bearing mice, we observed dramatic differences between NSG and C57BL/6 mice, not only of T and B cells but also other immune cells, including decreased monocytes, dendritic cells, and eosinophils. In immunocompetent brain tumor-bearing mice, we observed significant differences in the numbers of immune cells compared to non-tumour-bearing controls, primarily in the brain; however, significant differences were also in the spleen, lymph node and bone marrow, particularly in the subsets of monocytes, T cells and dendritic cells. Sub-lethal irradiation caused lasting changes in immune composition.

**Conclusions:** Our quantitative characterization of different immune compartments in orthotopic murine models of glioblastoma may allow for a better-informed selection of tumor models that are appropriate for the pre-clinical investigation of immunotherapies for glioblastoma.

## Introduction

A diagnosis of glioblastoma comes with a poor prognosis and patient survival of 7-17 months^1^. There have been no life-prolonging advances in systemic therapies beyond the standard of care in the last 20 years^2^ despite research activity that has been well-reviewed elsewhere^3^. Recently, immune-based therapies offer an exciting new approach for treating glioblastoma. Clinical trials activity includes immune checkpoint inhibitors, oncolytic viruses, autologous vaccines, and chimeric antigen receptor (CAR) T cells but so far there have been no durable survival benefits^4,5^, and it has been proposed that the immune-suppressive microenvironment of glioblastoma is a significant hurdle. One key challenge for developing new immunotherapies is finding suitable animal models to test effects of treatment on both targets and bystander cells. While developing GD2-targeted immunotherapies for brain tumors^6^ we identified a lack of detailed information on the immune compartments in pre-clinical brain tumor models. This gap in knowledge limits the ability of researchers to select the best model for investigating new immune-based treatments for brain tumors.

The different mouse models of brain tumors, including those generated from CT2A, GL261 and SB28 cells are well-characterized elsewhere^7^. Yet, although this publication provides a detailed description of the advantages and disadvantages of each model, there is no characterization of the immune compartments in these mice. Others have performed a detailed characterization of the changes in the tumor-localized immune cells present within CT2A and GL261 syngraft models, however, changes in systemic immune compartments such as spleen and bone-marrow were not characterized ^8^. This is despite the fact that glioblastoma is now acknowledged to have profound effects on the systemic immune response^5,9–11^. To the best of our knowledge, there is no current literature describing the baseline state of the whole immune system in these syngeneic models.

The immune-deficient NOD-SCID gamma null (NSG) mouse strain has been engineered to lack T, B and NK cells and together with a SIRPA polymorphism enables the successful engraftment of human cells, and is frequently used to model human gliomas^6^. These mice are highly immune-compromised with defects beyond a lack of adaptive immune cells: Interleukin-2Rγ (IL-2Rγ) allelic mutations and the NOD genetic background impair lymph node (LN) organogenesis^12^ and the function of dendritic cells (DCs) and macrophages^13,14^. As with the syngenic models, there has been very limited investigation of the immune system in NSG mice with engrafted gliomas.

To advance development of immunotherapies for glioblastoma we have undertaken to fully define the baseline immune system in mice with brain tumours. Using three immunocompetent models in C57BL/6 mice and two patient-derived xenograft models in NSG mice, we have applied high-parameter flow cytometry to systematically characterize the differences in immune cells in brain, spleen, cervical lymph nodes (LNs), and bone marrow (BM). In the course of this study we have also investigated the impact of reporter gene expression on immunogenicity, due to reports that these transgenes can be immunogenic in some strains of mice^15–17^, and characterised the impact of whole body irradiation, which is frequently used to allow adoptive cell transfer in immune-competent models. Together, our findings provide important new knowledge of the baseline immune state in mouse brain tumor models derived from cell lines widely used in the field. This knowledge can be leveraged in future studies to select the most appropriate system for testing novel immunotherapies.

## Results

We used a modified high-parameter flow cytometry panel, based on that described by Liu et al.^18^, to identify the immune cells in mice with and without orthotopic brain tumors^18^. In contrast to the individual detailed flow cytometry panels provided by Liu et al., we found that with careful consideration of the available fluorophores we could combine the individual panels into a single panel for analysis. We found this allowed for the identification of lymphocytes, granulocytes and monocytes (Fig 1A), dendritic cells (Fig 1B) and macrophages (Fig 1C) in the spleens, LNs, BM, and brains of the mice, as well as of microglia in the brain (Fig 1D). Microglia are defined by a CD45^int^ CD11b^+^ F4/80^+^ Ly-6C^-^ phenotype using this panel, and are brain resident cells derived during embryonic development. Microglia are thus distinct from infiltrating BM-derived myeloid lineages although BM-derived cells that have taken on a microglia-like phenotype can also be captured by these markers^19–21^. In addition, we have used a counting bead approach to enable reporting of both total numbers and proportions of immune cell types.

**Figure 1.**
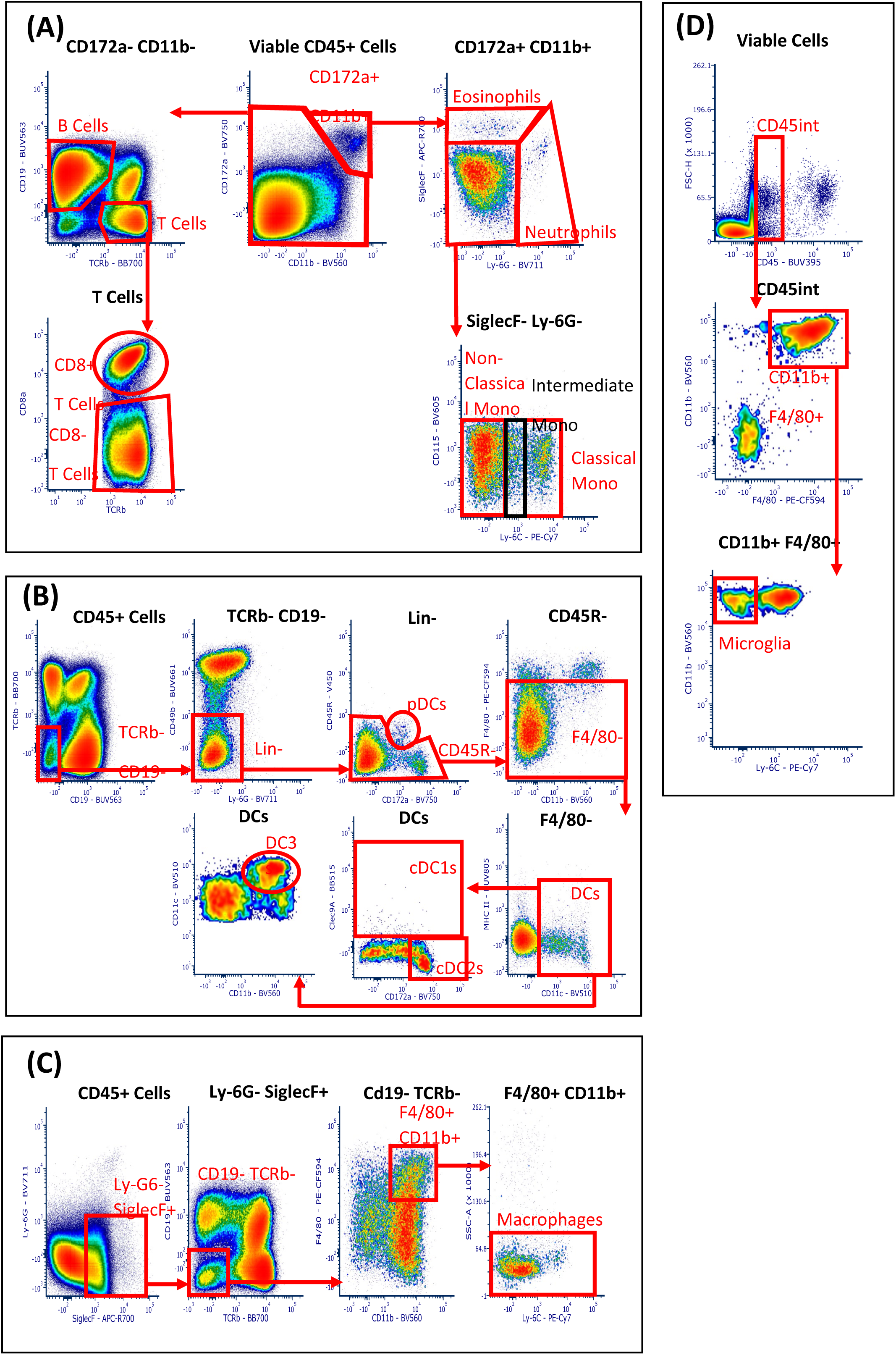
Indicative gating strategy to identify immune cells in the brain of a healthy C57BL/6 mouse. (A) Lymphocyte, monocyte and granulocyte gating, (B) Dendritic cell gating, (C) Macrophage gating, (D) Microglia gating.

### Enumeration of immune cell types in non-tumor-bearing C57BL/6 and NSG mice

First, we compared the immune compartments in sex and age-matched non-tumor-bearing C57BL/6 and NSG mice. As expected, we identified no B or T cells in any tissue of NSG mice (Fig 2A-D). In the brains, we observed a significant reduction in the number of eosinophils and a significant increase in the microglia in NSG mice compared to C57BL/6 mice (Fig 2A). In the spleen, we observed a significant increase in the macrophages in NSG mice compared to C57BL/6 mice, and a significant decrease in the monocytes, DCs and eosinophils in NSG compared to C57BL/6 mice (Fig 2B). We observed a significant decrease in the monocytes, DCs, neutrophils, and eosinophils in the LNs of NSG mice compared to C57BL/6 mice (Fig 2C). Finally, in the BM, we observed a significant decrease in macrophages, DCs, and eosinophils in the NSG compared to C57BL/6 mice (Fig 2D). The numerical defects observed in the immune cells of NSG compared to C57BL/6 mice suggest that prior study of immune system-brain tumor interactions may have been more limited in NSG mice, even for cells of the myeloid compartment, which have often been considered as relatively intact in NSG mice^22^.

**Figure 2.**
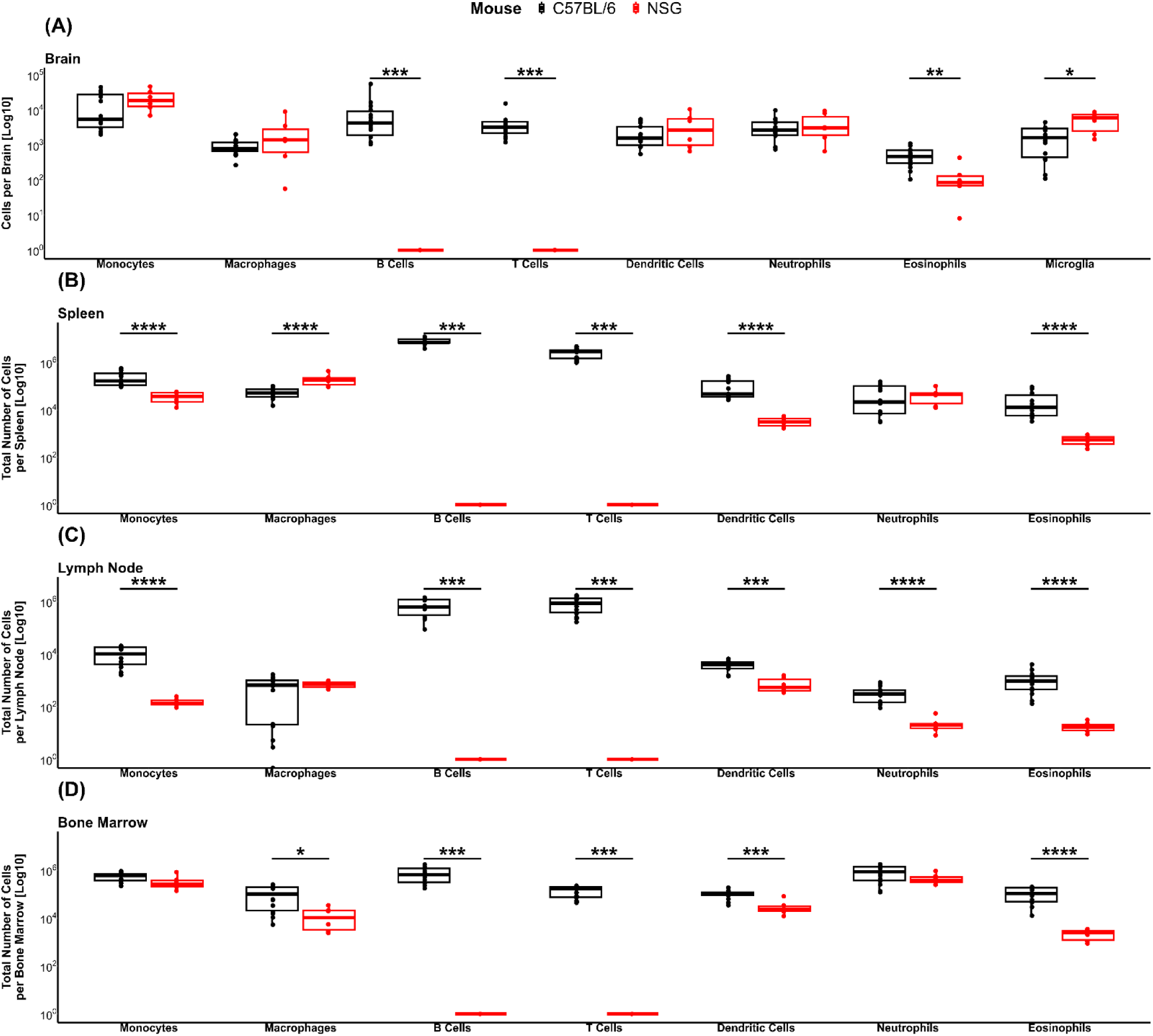
A comparison of immune cells in control C57BL/6 and NSG mice is shown as the total number per tissue. (A) Brain, (B) Spleen, (C) Lymph node, (D) Bone marrow. Data represent the median, second and third quartiles of n = 14 C57BL/6 mice and n = 6 NSG mice. P values *p<0.05, **P<0.01, ***p<0.001, ****p<0.0001.

### Immune cell frequency distribution in C57BL/6 and NSG mice with or without orthotopic brain tumors

Next, we investigated immune cells in the brain, spleen, LNs, and BM of non-tumor bearing (control) mice compared to mice with orthotopic brain tumors. Tumor cells were injected into the right cerebral hemisphere and allowed to grow for 3-4 weeks. All glioma cell lines were engineered to express the disialoganglioside GD2, (our target antigen of interest) by stable expression of murine GD2 and GD3 synthases. GD2 is conserved between mice and humans and is expressed at low level in normal brain of both species^6^. In immunocompetent C57BL/6 mice, we assessed the effect of intracranial tumors of GD2-expressing CT2A, CT2A-luc/GFP, GL261, GL261-luc/GFP, and SB28-luc/GFP glioma cell lines on the proportions of different leukocyte subsets. We compared the survival times of tumor-bearing mice with the parental CT2A, CT2A-luc/GFP, GD2-CT2A, and GD2-CT2A-luc/GFP cell lines from previous studies and observed no significant differences in mean survival (Fig S1).

Data is visually summarised in Figure 3A-D, with mean, standard error of the mean (SEM) and p values in Supplementary Table S1). As expected, the most dramatic differences in immune cell populations were observed in the brain. Mean, standard error of the mean (S.E.M) and p values are in Table S1. Specifically, we detected a significant increase in the proportion of monocytes in the brains of mice bearing CT2A, CT2A-luc/GFP and GL261 tumors compared to control mice, although unexpectedly, these cells were significantly reduced in the brains of mice bearing SB28-luc/GFP tumors, which instead had an increase in macrophages. Notably, BM-derived monocytes have been reported to support microglia in disease states and may be recruited before differentiation to supplement the brain resident microglia cells^19–21^.

**Figure 3.**
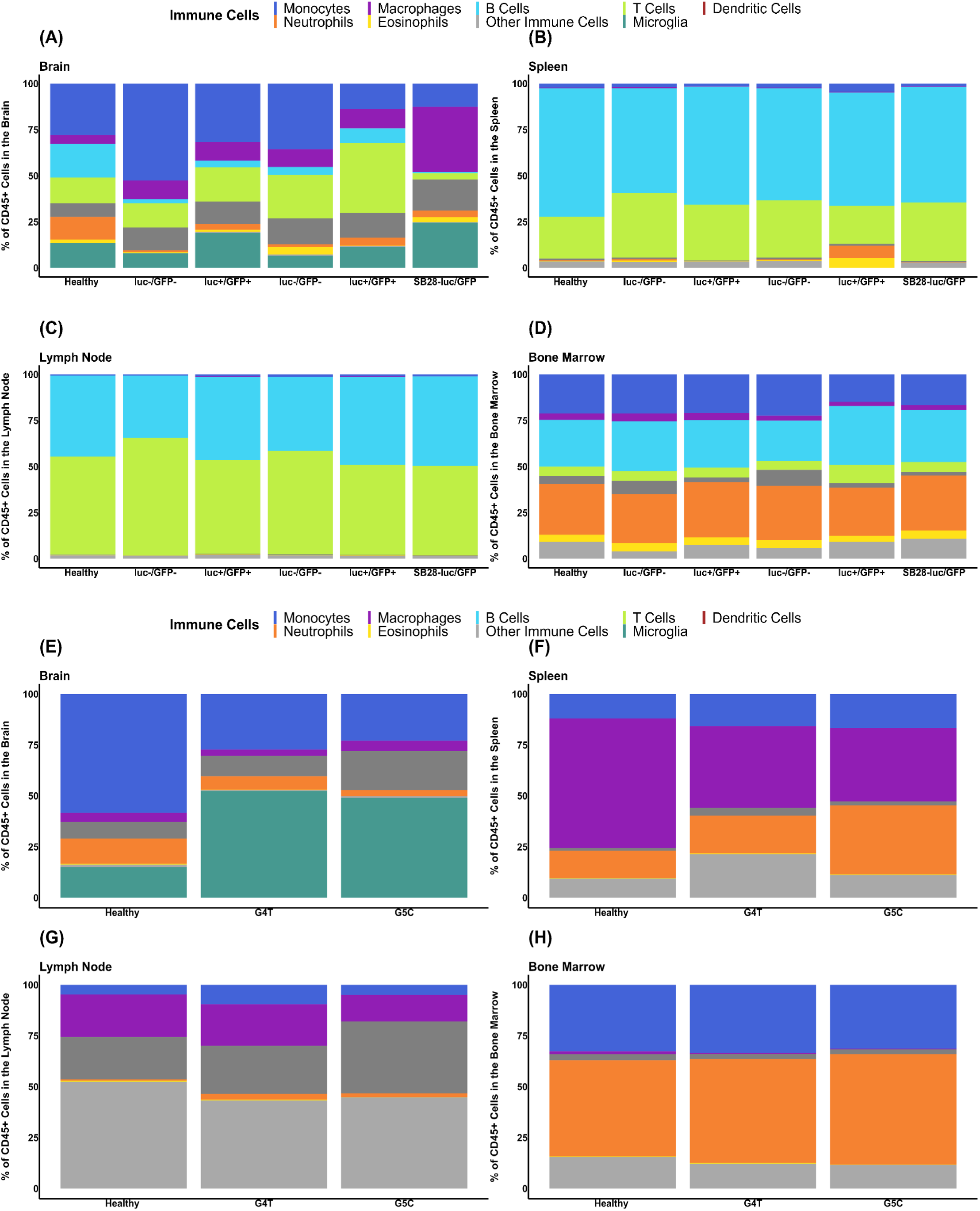
Immune cells as a percentage of the total CD45^+^ viable cells from (A) Brains of C57BL/6 mice, (B) Spleens of C57BL/6 mice, (C) Lymph nodes of C57BL/6 mice, (D) Bone marrow of C57BL/6 mice, (E) Brains of NSG mice, (F) Spleens of NSG mice, (G) Lymph nodes of NSG mice, (H) Bone marrow of NSG mice. Data represent the mean of n = 14 C57BL/6 mice, n= 11 CT2A tumor-bearing mice, n = 17 CT2A-luc/GFP tumor-bearing mice, n= 11 GL261 tumor-bearing mice, n = 5 GL261-luc/GFP tumor-bearing mice, n = 5 SB28-luc/GFP tumor-bearing mice, n = 6 NSG mice, n = 6 G4T tumor-bearing mice, and n = 6 G5C tumor-bearing mice. P values *p<0.05, **P<0.01, ***p<0.001, ****p<0.0001.

We observed a significant decrease in B cells in the brains of all tumor-bearing mice compared to the control mice. We also observed a significant increase in the percentage of dendritic cells in the brains of the CT2A-luc/GFP, GL261-luc/GFP, and SB28-luc/GFP tumor-bearing mice compared to control mice. Furthermore, we observed an increase in the percentage of T cells in the brains of CT2A-luc/GFP, and GL261-luc/GFP, and SB28-luc/GFP tumor-bearing mice compared to control mice (Fig 3A), with no apparent decrease in the spleen or increase in the BM of T cells in tumor-bearing mice despite previous reports^10^ (Fig 3A, B, D). Of note, we did not see significant changes in most immune cell types in the spleen of the tumor-bearing compared to control mice, apart from an increase in the proportion T cells in the CT2A tumor-bearing model, a reduction in the macrophages and dendritic cells in the CT2A-luc/GFP model and a reduction of dendritic cells in the SB28-luc/GFP tumor-bearing model (Fig 3B). There were also very few significant changes in the LNs or BM between tumor-bearing and control mice (Fig 3C-D).

In immunodeficient NSG mice, we assessed the effects of brain tumors generated from two different glioblastoma patient-derived glioma neural stem (GNS) cell lines, CCB-G4T-luc/RFP and CCB-G5C-luc/RFP^6,23^ (Fig 3E-H and Supplementary Table S1). We observed an increase in the frequency of microglia in the brains of mice bearing either of the patient-derived tumors compared to control mice, with a corresponding decrease in monocyte frequency (Fig 3E). However, one of the most notable differences observed was the increase in “other immune cells” in the spleen, LN and BM of NSG mice with or without a tumor compared to C57BL/6 mice; these are CD45^+^ cells that do not have other lineage markers and that we consider to be immature or defective cells^24^ (Fig 3F-H). Other subtle changes in immune cell populations were observed between mice bearing different tumors and were assessed quantitatively below as the absolute numbers of cells in these different immune cell subsets.

### Enumeration of immune cell types in C57BL/6 with orthotopic brain tumors with or without luciferase and GFP

As the immunogenic properties of luciferase-expressing cells have been reported as minimal in C57BL/6 mice^17^ and evident in other strains^15^, we first quantified differences in immune cell composition between the luciferase-expressing and non-expressing GL261 and CT2A cell lines implanted as orthotopic brain tumor models (Fig S2). To summarize these findings, we observed changes in immune cell number between tumor-bearing mice with and without reporter genes. Although the number of tumor-infiltrating cells in the brain (Fig S2A,E) suggested that the reporter molecules were not immunogenic, changes observed in the spleen of the CT2A tumor-bearing mice could indicate an increased immune response to reporter molecules in CT2A-luc/GFP tumor-bearing mice (Fig S2B) rather than in GL261 tumor-bearing mice (Fig S2F).

### Absolute numbers of immune cell types in C57BL/6 mice with or without orthotopic brain tumors

Although the analysis above revealed several changes in immune cell type frequencies between control and tumor-bearing mice, it is also important to consider potential changes in absolute numbers. Following our analysis of the differences between the tumors expressing luciferase/GFP and those not expressing luciferase/GFP, we have focused this analysis on luciferase-expressing tumor models because we believed that the value of longitudinal in vivo monitoring of tumor growth exceeded any potential detriment that immunogenic reporter molecules might have on immune cell composition.

Figure 4A shows the number of each immune cell type within brains of immunocompetent tumor-bearing and control mice. First, we observed a significant increase in the number of monocytes in brains of CT2A-luc/GFP tumor-bearing mice and SB28-luc/GFP tumor-bearing mice compared to control C57BL/6 mice, and a significant increase in macrophages in brains of all mice bearing syngeneic tumors compared to control mice. Second, we observed a significant increase in T cells in brains of the CT2A-luc/GFP and GL261-luc/GFP tumor-bearing mice compared to control mice. Interestingly, SB28-luc/GFP tumor-bearing mice were not observed to have the same increase in T cells. Third, DCs in the brains of SB28-luc/GFP and CT2A-luc/GFP tumor-bearing mice were also significantly increased compared to control mice, and eosinophils were significantly increased in the CT2A-luc/GFP tumor-bearing mice compared to control mice. Finally, we observed a significant increase in microglia in the CT2A-luc/GFP and SB28-luc/GFP tumor-bearing mice compared to control mice (Fig 4A).

**Figure 4.**
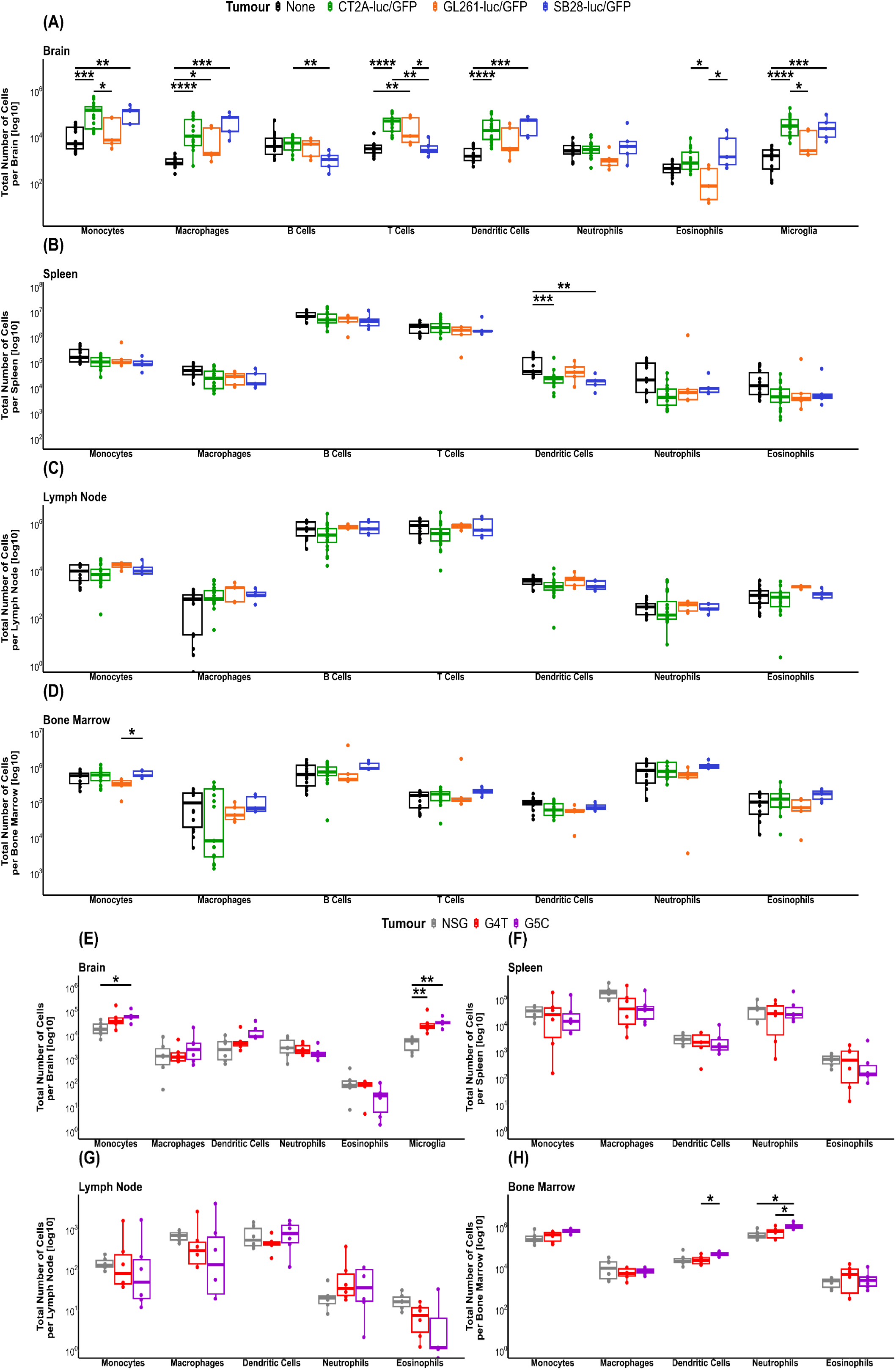
Absolute numbers of immune cell populations in control mice and mice with the indicated brain tumors. Graphs analyze immunocompetent C57BL/6 mice (A-D), or immunodeficient NSG mice (E-G). (A, E) Brains, (B, F) Spleens, (C, G) Lymph Nodes, (D, H) Bone marrow B cells and T cells are not shown for the NSG models as they are deficient in these cell types. Data represent the median, second and third quartiles of n = 14 C57BL/6 mice, n = 17 CT2A-luc/GFP tumor-bearing mice, n = 5 GL261-luc/GFP tumor-bearing mice, n = 5 SB28-luc/GFP tumor-bearing mice, n = 6 NSG mice, n = 6 G4T tumor-bearing mice, and n = 6 G5C tumor-bearing mice. P values *p<0.05, **P<0.01, ***p<0.001, ****p<0.0001.

When considering the systemic immune changes, we observed a significant decrease in splenic DCs of CT2A-luc/GFP and SB28-luc/GFP tumor-bearing mice compared to control mice and a significant decrease in splenic neutrophils and eosinophils in CT2A-luc/GFP tumor-bearing mice compared to control mice (Fig 4B). Remarkably, we observed no significant changes in any of the immune cell subsets in LNs or BM in the immunocompetent models compared to control mice (Fig 4C-D).

In summary, these analyses of immunocompetent brain tumor models reveal a significant increase in brain infiltrating immune cells in CT2A-luc/GFP tumor-bearing bearing mice, and to a lesser extent in SB28-luc/GFP tumor-bearing mice compared to the control mice while spleens of tumor-bearing mice show a trend toward fewer immune cells than in control mice.

### Absolute numbers of immune cell types in NSG mice with or without orthotopic brain tumors

Next, we investigated the immunodeficient models. In the brain, we observed a significant increase in total monocytes in G5C-luc/RFP tumor-bearing mice compared to control mice and a significant increase in microglia in all tumor-bearing mice compared to control NSG mice (Fig 4E), which is consistent with our findings in immunocompetent models. In contrast, we observed no significant changes in spleen or LNs of immunodeficient models (Fig 4F-G). Finally, we observed a significant increase of neutrophils in BM of G5C-luc/RFP tumor-bearing mice compared to control mice and no differences in any other cell type (Fig 4H). Overall, the presence of a brain tumor had little effect on immune cell populations in NSG mice.

### Enumeration of T cell, monocyte and DC subsets in brain tumor models

Given the diversity that exists within immune cell types, we next investigated the subsets of immune cells in greater detail. We performed sub-analyses of monocytes to enumerate classical phagocytic monocytes, non-classical inflammatory antigen cross-presenting monocytes, and transitional intermediate monocytes^25^. We also subdivided T cells into CD8^+^ and CD8^-^ subsets, and differentiated between anti-tumor cross-presenting classical dendritic cells type 1 (cDC1s), pro-tumor classical dendritic cells type 2 (cDC2s), and inflammatory dendritic cells type 3 (DC3s).

In considering the brains of tumor-bearing immunocompetent mice (Fig 5A), we observed a significant increase in the number of classical and non-classical monocytes, but not intermediate monocytes, in CT2A-luc/GFP and SB28-luc/GFP tumor-bearing mice compared to control C57BL/6 mice. When analysing CD8^+^ and CD8^-^ T cells separately, we observed the same trends as for bulk T cells, where compared to control mice, both subsets were increased in the CT2A-luc/GFP and GL261-luc/GFP models, but not in the SB28-luc/GFP model. In DC subsets, we observed a significant increase in the cDC2s and DC3s in CT2A-luc/GFP and SB28-luc/GFP mice compared to control C57BL6 mice, whereas cDC1 numbers were unchanged.

**Figure 5.**
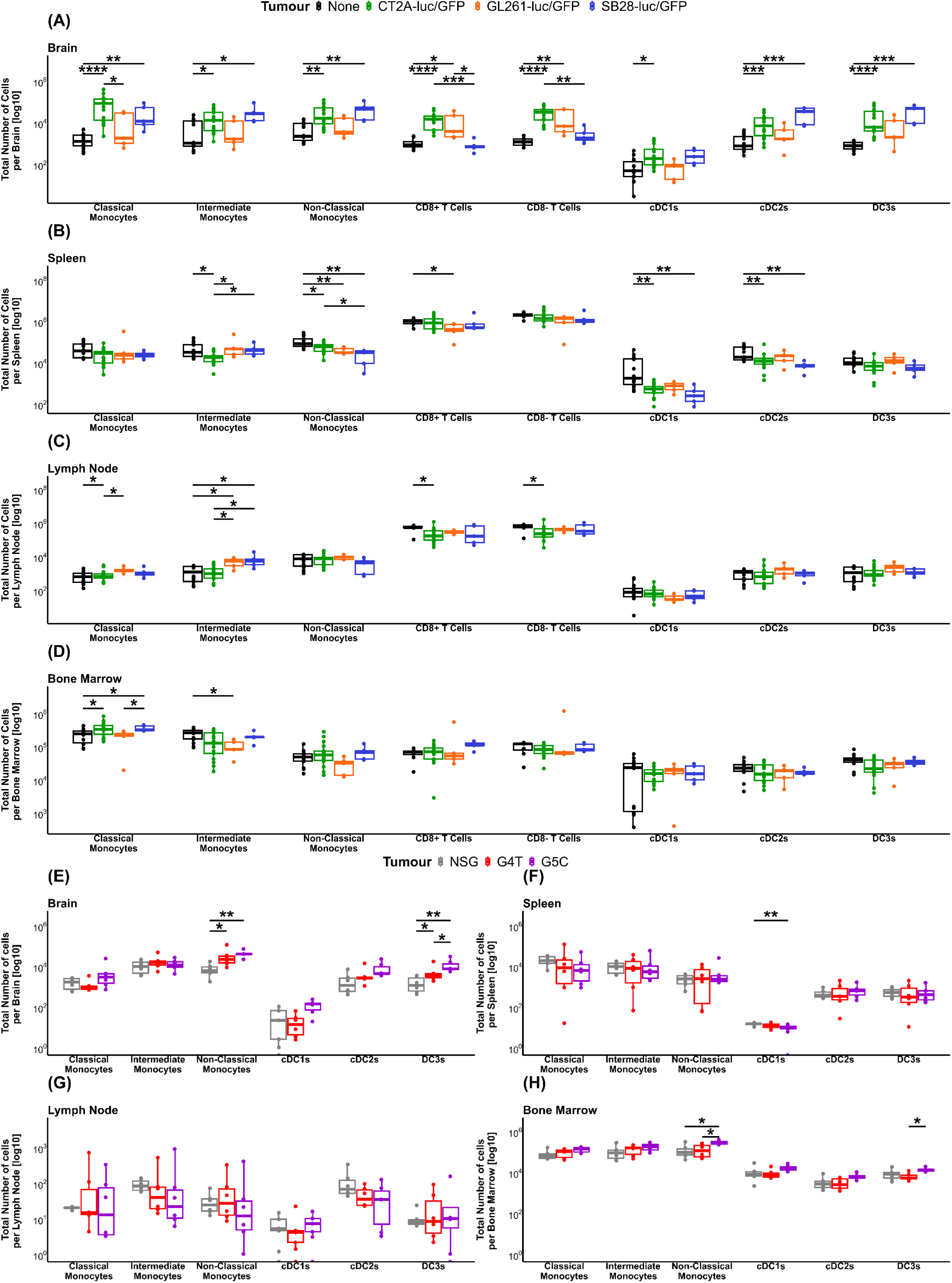
Absolute numbers of monocyte, T cell and DC subsets in control mice and mice with the indicated brain tumors. Graphs analyze immunocompetent C57BL/6 mice (A—D), or immunodeficient NSG mice (E-G). (A, E) Brains, (B, F) Spleens, (C, G) Lymph Nodes, (D, H) Bone marrow B cells and T cells are not shown for the NSG models as they are deficient in these cell types. Data represent the median, second and third quartiles of n = 14 C57BL/6 mice, n = 17 CT2A-luc/GFP tumor-bearing mice, n = 5 GL261-luc/GFP tumor-bearing mice, n = 5 SB28-luc/GFP tumor-bearing mice, n = 6 NSG mice, n = 6 G4T tumor-bearing mice, and n = 6 G5C tumor-bearing mice. P values *p<0.05, **P<0.01, ***p<0.001, ****p<0.0001

Many changes in these immune cell subsets were also observed elswhere in the immune system. In the spleen (Fig 5B), we observed a significant decrease in intermediate monocytes in the CT2A-luc-GFP tumor-bearing mice compared to control C57BL/6 mice, and a significant decrease in non-classical monocytes in all tumor-bearing mice compared to control C57BL/6 mice. There were very few changes to T cell subsets, with the only difference being a significant decrease in CD8^+^ T cells in GL261-luc/GFP tumor-bearing mice compared to control C57BL/6 mice. For DCs, we observed a significant decrease in splenic cDC1s and cDC2s in SB28-lucGFP mice, while there was only a significant decrease in cDC1s in CT2A-luc/GFP tumor-bearing mice compared to control C57BL/6 mice. Within LNs, we observed a significant increase in the intermediate monocytes in GL261-luc/GFP and SB28-luc/GFP tumor-bearing mice compared to control C57BL/6 mice, and a significant decrease in both CD8^+^ and CD8^-^ T cells in CT2A-luc/GFP tumor bearing mice compared to control C57BL/6 mice (Fig 5C). Finally, we examined the BM (Fig 5D), where we observed a significant increase in classical monocytes in CT2A-luc/GFP and SB28-luc/GFP tumor-bearing mice and a significant decrease in intermediate monocytes in GL261-luc/GFP mice compared to control C57BL/6 mice.

We then investigated changes to leukocyte subsets in the immunodeficient models. In the brain, we observed a significant increase in non-classical monocytes and DC3s in both tumor-bearing models compared to the control NSG mice (Fig 5E). In contrast, no significant changes were observed in spleen or LNs (Fig 5F-G), whereas we observed a significant increase of non-classical monocytes in BM of G5C-luc/GFP mice compared to the control NSG mice (Fig 5H).

Together, these detailed analyses reveal that most changes in immune cell subsets occur in brain and spleen of tumor-bearing mice, with minimal changes in the composition of immune cell subsets in LNs and BM. Meanwhile, the NSG mice showed very few changes in extracranial immune cell subsets in the tumor-bearing mice compared to control mice.

### Immunofluorescence Staining of Mouse Brains

To confirm the presence and location of immune cells in the brains of the mice, we conducted immunofluorescence staining for CD3 and CD11b (Fig 6). Representative images of the staining include a control mouse brain (Fig 6A), and brains of mice bearing GFP-expressing tumors (Fig 6B). The fluorescence intensity stained for each antibody was quantified separately for tumor and non-tumor areas using FIJI (Fig 6C-D). In all models CD3+ cells were detected at low levels, and we observed no significant difference in the intensity of the CD3 staining in the tumor region of the brains of the tumor-bearing mice compared to the non-tumor area. There was also no significant difference in the CD3 intensity in brains of tumor-bearing mice compared to control mice (Fig 6C). In contrast, we observed a significant increase in the fluorescence intensity of CD11b staining only in the tumor areas of all tumor-bearing mice compared to the non-tumor areas, which in turn did not display significant differences compared to brains of the control mice (Fig 6D).

**Figure 6.**
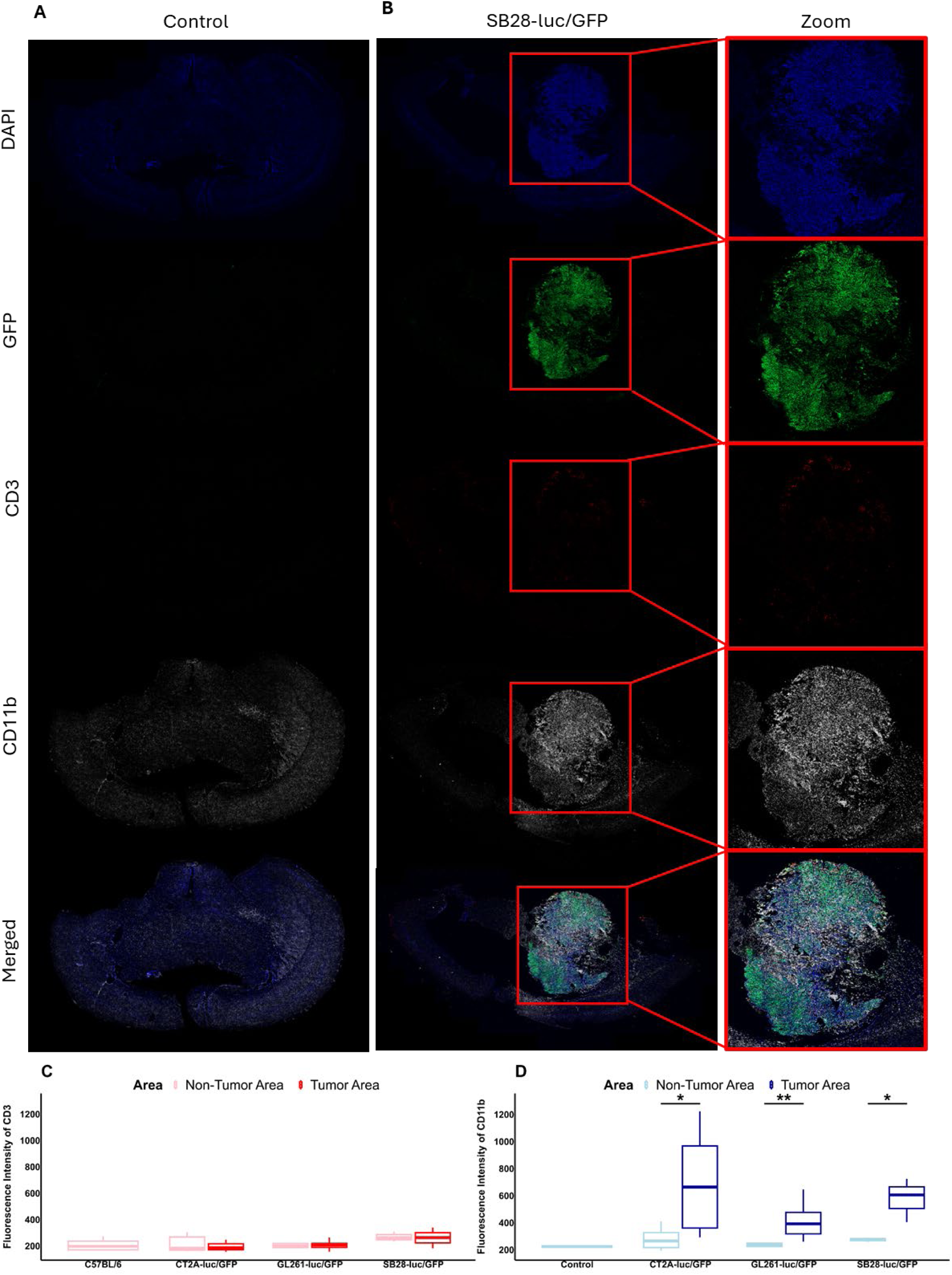
Representative immunofluorescence staining of C57BL/6 mouse brain sections to identify CD3^+^ T cells and CD11b^+^ cells. Panels show from top to bottom DAPI, GFP, CD3, CD11b, and merged channels. (A) Control C57BL/6 mouse, (B) representative image of the brain of a GFP tumor-bearing mouse, (SB28-luc/GFP) tumor-bearing mouse, (C) Quantitative data of immunofluorescence intensity of CD3 staining (D) Quantitative data of immunofluorescence intensity of CD11b staining.

### Effect of whole body irradiation on immune cell types

We then considered the effects of sub-lethal whole-body x-ray irradiation (WBI), which includes the brain, on the CT2A-luc/GFP tumor-bearing mice. Sub-lethal irradiation is frequently used to lymphodeplete mice and promotes engraftment of adoptively transferred cells such as CAR-T cells^26,27^. Irradiation is also reported to, at least initially, reduce immunosuppression by depleting pro-tumor immune cells such as tumor-associated macrophages as well as increase tumor immunogenicity both by upregulating expression of damage-associated molecular pattern molecules and inducing tumor cell death that results in release of and increased presentation of tumor antigens^28^. Despite routine use in adoptive T cell transfer studies, the baseline effects of WBI have not been well characterised. In contrast to adoptive cell transfer studies that often use doses of 5 Gy^29^ or 6 Gy^30^, studies of the effects of whole-body irradiation on immune cell composition were limited to dose ranges up to 1 Gy (with analysis of spleen and BM)^31^ or up to 2 Gy (with analysis only of splenocytes)^32^.

To evaluate the effects of WBI on non-tumor-bearing mice, we irradiated C57BL/6 mice with a range of X-ray doses from 0.5 Gy to 5 Gy and analyzed immune cells 5 days later. For initial analyses, we enumerated bulk CD45^+^ immune cells in spleen, peripheral blood and BM (Fig 7A) and, compared to non-irradiated controlmice, we observed significant decreases in the spleen at all WBI doses, with the most marked decrease seen at the 5 Gy dose. Although we observed no significant changes in peripheral blood numbers of CD45^+^ cells at any WBI dose compared to control mice, there was a clear trend toward reduced numbers of circulating CD45^+^ immune cells at higher WBI doses. We also observed a significant reduction in CD45^+^ immune cells in BM of mice treated with 5 Gy WBI compared to all other mice.

**Figure 7.**
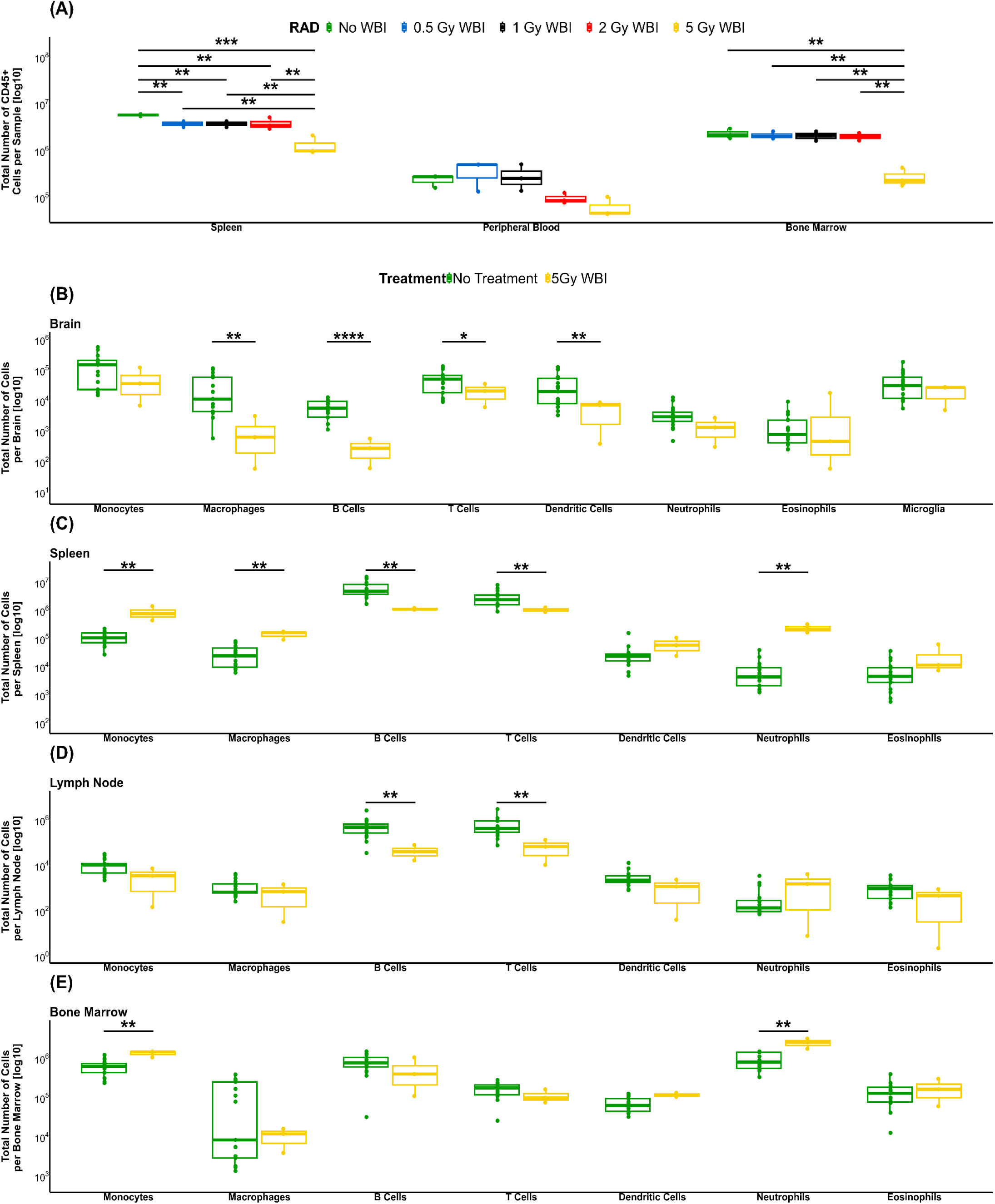
Immune cells in C57BL/6 mice with and without whole body x-ray irradiation. (A) dose comparison in tumor-free mice, analyzing the total number of CD45^+^ leukocytes in the spleen, peripheral blood and bone marrow 5 days post-irradiation, (B) immune cells in the brains of CT2A-luc/GFP tumor-bearing mice with and without WBI analyzed at humane endpoint, (C) immune cells in the spleens of CT2A-luc/GFP tumor-bearing mice with and without WBI analyzed at humane endpoint, (D) immune cells in the lymph nodes of CT2A-luc/GFP tumor-bearing mice with and without WBI analyzed at humane endpoint, (E) immune cells in the bone marrow of CT2A-luc/GFP tumor-bearing mice with and without WBI analyzed at humane endpoint. Mean survival = 22.2 days without WBI vs 28.5 days with WBI. Data represent the median, second and third quartiles of n = 3 per group for non-tumor-bearing irradiated mice, n = 17 CT2A-luc/GFP tumor-bearing mice, and n = 3 5Gy WBI CT2A-luc/GFP tumor-bearing mice. P values *p<0.05, **P<0.01, ***p<0.001, ****p<0.0001.

Next, we treated CT2A-luc/GFP tumor-bearing mice with 5 Gy WBI 7 days after tumor cell implantation. At the humane endpoint (approximately four weeks after tumor cell implantation), we compared the immune cells in these irradiated mice to CT2A-luc/GFP tumor-bearing mice that did not receive WBI. In the brain, we observed a significant reduction in macrophages, B cells, DCs, and neutrophils when tumor-bearing mice received 5 Gy WBI (Fig 7B). In the spleen (Fig 7C), we observed a significant reduction in B cells and T cells but a significant increase in monocytes, macrophages, neutrophils and eosinophils in WBI-treated mice compared to non-irradiated tumor-bearing mice. In the LNs (Fig 7D), we again observed a significant reduction in B cells and T cells following irradiation. Finally, in the BM, we observed a significant increase in monocytes and neutrophils in the WBI-treated mice compared to the non-irradiated tumor-bearing mice, while all other immune cells were unaffected by WBI (Fig 7E). Taken together, this would suggest that most immune cells have returned to normal levels post-WBI at the humane endpoint. Two exceptions to this trend include the tumor-bearing brains, where many immune cell populations were still altered following WBI, and in B cells and T cells, which seem to take longer to repopulate spleen and LNs than other immune cells.

## Discussion

Although the immunotherapy revolution has transformed the treatment paradigm for many cancers, it is not standard treatment for glioblastoma patients and remains an active area of research^5,33,34^. Immunotherapy approaches such as immune-checkpoint inhibitors may be useful in < 3% of glioblastoma patients, and although CAR-T cell therapy may have broader application, early-phase clinical trial data are limited^35–37^. Further research is required to enable immune therapies to combat features of the glioblastoma tumor microenvironment that limit anti-tumor immune activity including antigen heterogeneity and a phalanx of immunosuppressive molecules and cell types, particularly those of myeloid origin^38,39^. Moreover, in glioblastoma, immune suppression extends system-wide and includes DC dysfunction, lymphopenia, T-cell sequestration and elevated circulating myeloid-derived suppressor cells (MDSCs)^10,40–45^. To develop effective immune-based treatments it is therefore essential that we have appropriate pre-clinical models that recapitulate these features of glioblastoma. Since we found little information about system-wide immune compartments in commonly used orthotopic models of murine brain tumor we have undertaken detailed studies to examine immune compartments in control NSG and C57BL/6 mice and in the same strains of mice hosting intracranial brain tumors.

One aspect of our studies focused on better defining the immune system of NSG mice beyond the expected loss of T, B and NK cells. Previous reports have described immature macrophages and dysfunctional DCs^13,14^. We now build on these previous findings by revealing significant changes in the absolute numbers of several immune cell types in multiple organs, when comparing NSG mice to immune competent C57BL/6 mice. In particular, we identified that monocytes and DCs were significantly reduced in the spleens and LNs of NSG mice, and eosinophils were significantly reduced in all tissues analyzed, compared to C5BL/6 mice. Interestingly, these systemic reductions in monocytes and DCs did not extend to the brain where, compared to C57BL/6 mice, we found increased numbers of microglia and no changes to monocyte or DC numbers in NSG mice.

As expected, the implantation of brain tumors in C57BL/6 mice resulted in significant changes in brain immune microenvironment, with a significant increase in classical and non-classical monocytes, macrophages, CD8^+^ and CD8^-^ T cells, cDC2s, DC3s, and microglia in the brains of the CT2A-luc/GFP tumor-bearing mice compared to control mice. Further, we observed the same significant increases with the exception of the T cells in the brains of the SB28-luc/GFP tumor-bearing mice compared to control mice. This suggests the SB28-luc/GFP model may best represent the low T cell infiltration observed in human glioblastoma^46^. In immunodeficient NSG mice bearing orthotopic xenografts of patient-derived short-passage glioma neural stem cell lines, we observed significant increases in populations of non-classical monocytes and microglia in the brains of our tumor-bearing models compared to control mice.

The immunofluorescence staining of the brains revealed an increase in the CD11b staining that was consistent with the increase in monocytes and macrophages detected by flow cytometry in the tumor-bearing mice, as compared to the control mice. In contrast, the CD3 immunofluorescence staining did not show the same increase that was seen in the flow cytometry data of tumor-bearing mice compared to the control mice, likely due to the lower sensitivity of this assay compared to flow cytometry.

Given the intriguing findings of glioblastoma-mediated global immune suppression by others^10,40–45^, we investigated lymphoid organs such as the spleen and BM of tumor-bearing mice. In immunocompetent C57BL/6 mice, we found that the presence of a brain tumor induced few changes in the BM, and we did not observe the previously reported^10^ sequestration of T cells in this site. We also observed no changes in LNs. In the spleens of tumor-bearing mice, DCs, neutrophils and eosinophils were reduced, but no changes were observed in other populations. Our study, therefore, supports the concept that brain tumors induce some level of systemic immune suppression in mice, but this is not as profound or widespread as previously reported. Furthermore, this effect was not observed in NSG models, where the presence of a brain tumor did not lead to systemic reductions in any immune cell types beyond the immune defects already present within this strain. Limitations of our studies in immunocompetent orthotopic brain models include the use only of syngraft tumors, which are acknowledged to be less similar biologically to human glioblastoma than genetically engineered models^7^, and not examining the functional status of the enumerated immune cells, which remains an area for future investigation.

Our syngeneic models also overexpress the glioblastoma antigen GD2, which is ubiquitously expressed on the GNS lines^6^. GD2 is an important immunotherapeutic target under investigation for the treatment of glioblastoma^6,47–50^. GD2 is a glycosphingolipid which is identical in mice and humans, allowing toxicity testing of human-GD2-targeting therapeutics in mice^51^, and as such, is not expected to contribute to the immunogenicity profile. Although this was not formally tested in this study, we did not observe significant differences in mean survival times for CT2A models with and without GD2 or luc/GFP, although GD2-CT2A-luc/GFP has a longer survival tail.

Another new finding in this study was the significant effects of WBI on the immune system of C57BL/6 mice. WBI is used routinely in syngraft models to achieve CAR-T engraftment^29,30^, yet the global immune effects are rarely considered. We found that at the humane endpoint, approximately 4 weeks after irradiation, there was still a significant reduction in B cells and T cells in brain, spleen, and LNs of irradiated mice. We also observed a significant increase in monocytes, macrophages, neutrophils, and eosinophils in spleen of WBI mice. Importantly, we have shown that use of WBI in these models has effects beyond transient lymphodepletion and, in fact, in our study, there appears to be broad remodelling of the immune system following WBI, which will have consequences for any immune-targeted treatment.

This study highlights the importance of a broad assessment of immune cell numbers when deciding which pre-clinical model is appropriate for testing immune-based therapies. NSG mice are routinely used for xenograft models of CAR-T therapy because they enable human tumor and human T cells to be assessed. However, as we have shown, there are extensive differences in system-wide immune cell composition beyond the absence of B and T cells, and including a broad reduction in DCs, monocytes and eosinophils. Therefore, it will be difficult to adequately assess the endogenous immune effects of approaches that rely on systemic immune engagement, such as cytokine-enabled CAR-T^6,52^ or other combination therapy approaches^53–55^.

However, we did observe that system-wide reductions in the number of monocytes and DCs did not apply in the brain. These data suggest that, despite numerical and functional impairments at steady state, myeloid cell populations in NSG mice are responsive to the emergency demands imposed by the pathology of a brain tumor. Consequently, these data suggest that NSG brain tumor models may be suitable for studies of immunotherapies targeting myeloid cells, and for investigating interactions between CAR-T cell therapies and the local brain tumor immune microenvironment, which is typically dominated by macrophages and microglia. Nevertheless, further studies are needed to better define the function of these brain-resident myeloid cells in NSG mice.

In conclusion, this study presents a global overview of immune cell numbers in commonly used pre-clinical models of glioblastoma. In considering immunocompetent syngraft models, all three models had evidence of immunogenic tumors with extensive immune cell infiltration of the brain tumor, which is not reported for human glioblastoma and without the evidence of extensive systemic immune suppression described by others^10^. The SB28 model did appear most closely to represent human glioblastoma because the levels of T cells in the brain are the lowest of all the tumor models we investigated. Together, these findings will enable researchers in the field of glioblastoma immunotherapy to better interpret their results and we hope will support further development of immunotherapy for this devastating disease.

## Materials and Methods

### Patient-Derived Glioblastoma Cell Lines

The patient-derived GNS cell lines were generated as previously described ^56^. Briefly, an aliquot of dissociated glioblastoma cells was resuspended in 3 mL StemPro NSC medium (Thermo Fisher) and transferred to a T-25 tissue culture flask coated for 30 min at 37°C with Matrigel (Corning Incorporated, Corning, NY, USA) diluted 1/100 in PBS. The following day, the flask was rinsed twice with warm DMEM to remove debris and dead cells, and fresh StemPro NSC was added. Cell culture details are in the supplementary methods.

### C57BL/6 Syngenic Brain Tumor Cell Lines

Parental lines CT2A (Merck) GL261 (Division of Cancer Treatment and Diagnosis Tumour Repository, NCI) SB28-luc/GFP (DSMZ; Leibniz-InstitutDSMZ Deutsche Sammlung von Mikroorganismen und Zellkulturen GmbH) were engineered to express the GD2 antigen by transduction with ecotropic retrovirus encoding murine GD2 and GD2 synthases (MP9956:SFG.GD3synthase-2A-GD2synthase was a gift from Martin Pule Addgene plasmid # 75013 ; http://n2t.net/addgene:75013^57^; RRID:Addgene_75013; retrovirus generated from the Phoenix-Eco (ATCC Phoenix-ECO CRL-3214) packaging line), with two-rounds of FACS sorting to achieve a >99% GD2+ line. In addition, GD2-CT2A and GD2-GL261 were rendered GFP/F-luciferase positive via lentiviral transduction using pBliv MSCV-copGFP T2A fLuc and FACS sorting on GFP+GD2+ cells. Culture details are in the supplementary methods.

### Orthotopic Brain Tumor Models

Animal experiments were conducted under a protocol approved by the University of South Australia Animal Ethics Committee (#32-22 & #U08-23). A Stoelting motorized stereotactic alignment and injection unit was used to deliver 2 µL of tumor cells to the brain over a period of 4 minutes, 3 mm deep into the right hemisphere. Syngraft models were generated by the implantation of GD2-SB28-luc/GFP cells (5×10^3^ cells), GD2-CT2A cells (5×10^4^ cells), GD2-CT2A-luc/GFP cells (5×10^4^ cells), GD2-GL261 cells (1×10^5^ cells), and GD2-GL261-luc/GFP cells (1×10^5^ cells) into 6–8 weeks old female C57BL/6 mice (Australian BioResources, Moss Vale). Patient-derived cell lines G5C-luc/RFP (2×10^5^ cells) and G4T-luc/RFP (2 × 10^5^ cells) were injected into 6-8 weeks old female NSG mice (Ozgene, Perth). Mice received daily clinical checks and weekly bioluminescence imaging (BLI) for mice with luciferase expression to monitor tumor growth. BLI was performed using an IVIS Lumina S5 after intraperitoneal injection of 100 µL of 30 mg/mL luciferin in saline. Defined euthanasia endpoints included loss of >15% body weight from starting weight, a body condition score of <2 (under-conditioned)^58^, and neurological signs including head-tilt, loss of balance and circling movement.

### Mouse Whole-body Irradiation

Whole-body irradiation was conducted in X-Rad225 (Precision X-ray irradiation, Madison, CT). Each dose was calculated using an internal probe to deliver a total effective dose of 0.5 Gy, 1 Gy, 2 Gy, or 5 Gy. The non-tumor-bearing mice were analyzed 5 days post WBI. Analysis of the tumor-bearing mice and corresponding control mice was at the humane endpoint.

### Murine Tissue Samples

Spleens, bone marrow (from the left rear leg), cervical lymph nodes and brains were harvested from euthanized mice and mechanically dissociated utilizing 40 µm strainers in Roswell Park Memorial Institute (RPMI) 1640 Medium (ThermoFisher Scientific, Cat# 11875093) + 1% FBS (CellSera, Cat# AU-FBS/PG). Samples were centrifuged at 400g for 5 min and resuspended in FBS + 10% dimethyl sulfoxide (DMSO) (Sigma-Aldrich, Cat# D8418) and cryopreserved for later analysis. Alternatively, brains were processed for immunofluorescence analysis by fixation for 1 hour in 10% neutral buffered formalin, followed by overnight in 15% sucrose solution and overnight in 30% sucrose solution prior to embedding in optimal cutting temperature (OCT) compound.

### Flow Cytometry Staining and Analysis

Stained cells were analyzed on a BD FACSymphony A5 (BD Biosciences) with FCS Express v7 Flow Research Edition (De Novo Software, Pasadena, CA, USA). For further detail, see online supplemental methods; antibodies utilized are in Table S2.

### Immunofluorescence Staining and Analysis

Staining was conducted as previously described^6^. Briefly, sections were incubated with 14.6µg/mL rat anti-CD11b (M1/70.15; Novus Biologicals) and 4µg/mL rabbit anti-CD3 (Abcam) for 12 hours at 4°C. Sections were washed and then incubated with donkey anti-rat IgG 647 and goat anti-rabbit IgG 555 at a dilution of 1:500 for 1 hour at RT. For full details, see supplemental methods.

### Statistical Analysis

Data was graphed and analyzed using Rstudio Version 2023.06.1. The list of packages is in Table S3. Statistical analysis was conducted in Rstudio utilizing the Kruskal-Wallis test and the Pairwise Wilcoxon test with Benjamini & Hochberg multiple comparison post-tests. P values <0.05 were considered statistically significant.

## Acknowledgments

The authors acknowledge the support received from the NeuroSurgical Research Foundation (NRF). BG also acknowledges the support received through the NRF-Strong Enough to Live scholarship. MNT acknowledges the support of a Fay Fuller Foundation Fellowship. LME acknowledges the support of the Ray & Shirl Norman Cancer Research Trust and James & Diana Ramsay Foundation. Glioblastoma patient tissues were received from the South Australian Neurological Tumor Bank (SANTB) which is supported by Flinders University, Flinders Foundation and the NRF. The authors also acknowledge the assistance of the staff of the Core Animal Facility and the Veterinarian at the University of South Australia with the experiments. We acknowledge Steven Roberts (BD Biosciences) for technical advice in designing multicolor flow panels for use on the BD FACS Symphony. The authors also acknowledge the support of the Adelaide Health and BioMedical Precinct Cytometry Facility at SAHMRI. The Facility is generously supported by the Detmold Group, McMahon Family, Australian Cancer Research Foundation.This project was funded by Cancer Australia Priority-Driven Collaborative Cancer Research Scheme and The Kids’ Cancer Project (Grant ID 2020344).

## Data Availability

Data will be made available upon reasonable request.

## Author Information

Conceptualization (BG, TG, MPB, LME), Writing – original draft (BG), Writing – Review and Editing (BG, TG, BLG, MNT, SMP, MPB, LME, TG), Data curation (BG, PMK), Formal analysis (BG, EN, JK, BLG, MNT), Methodology (BG, BLG, MNT, SMP), Funding Acquisition (TG, LME, MPB, SMP), Supervision (TG, LME, MPB)

## Supplementary Information

Natureportfolio Reporting Summary

ARRIVE guidelines E10: author checklist

**Figure S1.**
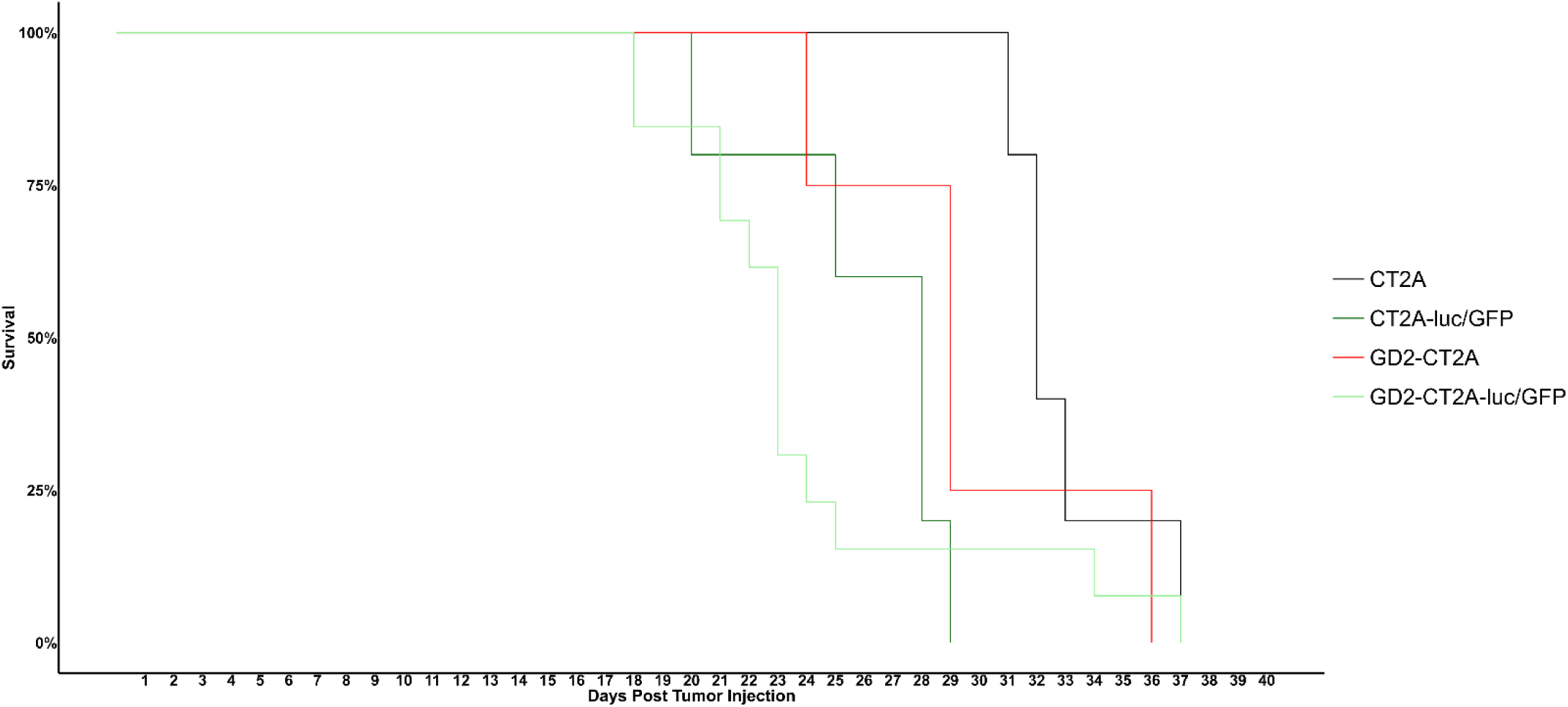
Survival time (to humane endpoint) for C57BL/6 tumour-bearing mice with (A) CT2A parental n=5 – median survival 32 days, CT2A-GFP/luc n=5 – median survival 28 days, GD2-CT2A n=4 – median survival 29 days, (B) GD2-CT2A-luc/GFP n=13 – median survival 23 days.

**Table S1.**
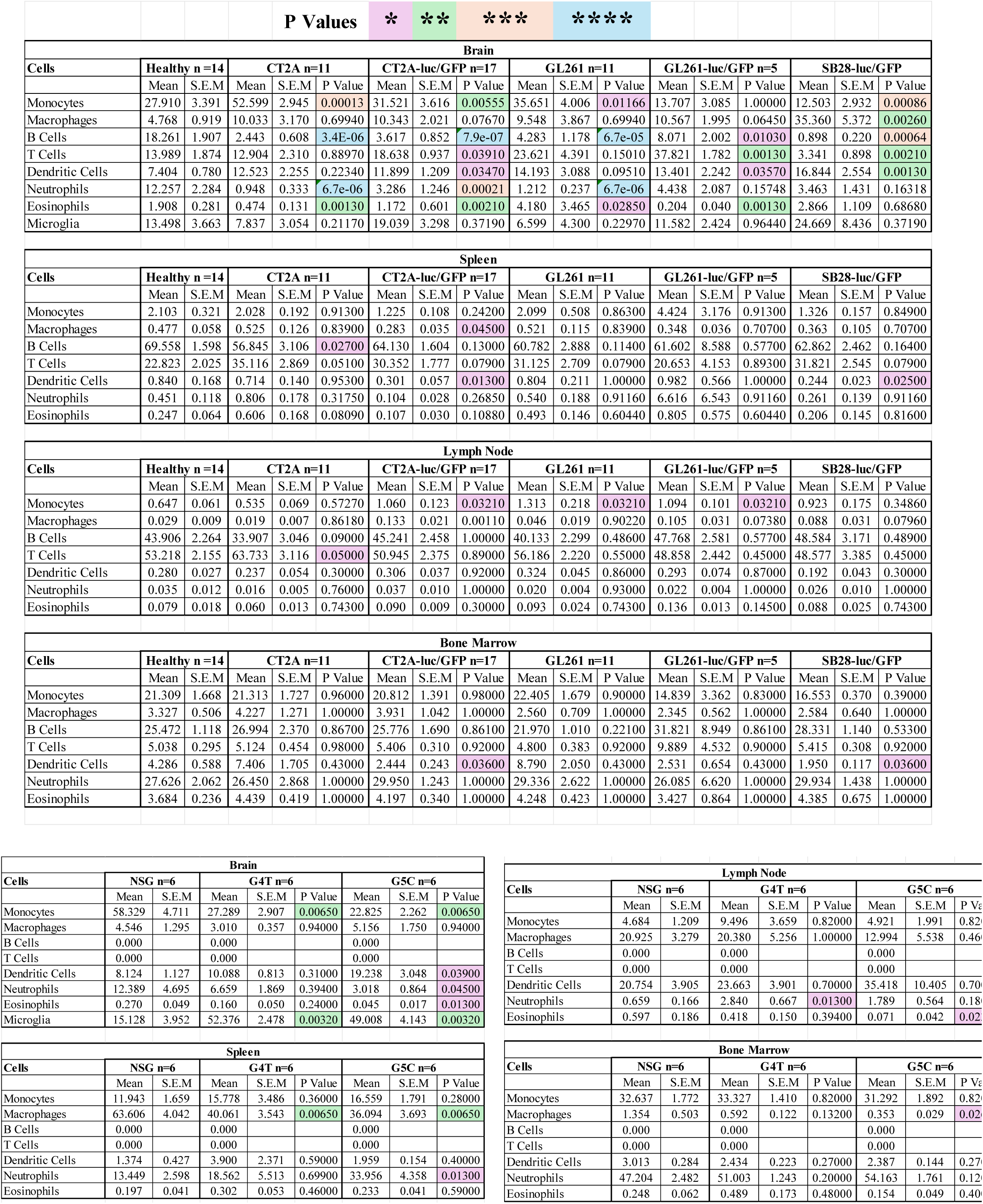
Percentage of CD45^+^ immune cells in mice with brain tumors in C57BL/6 and NSG mice. Data shows mean, SEM and p values corresponding to the data presented in Figure 3. P values *p<0.05, **P<0.01, ***p<0.001, ****p<0.0001.

**Figure S2.**
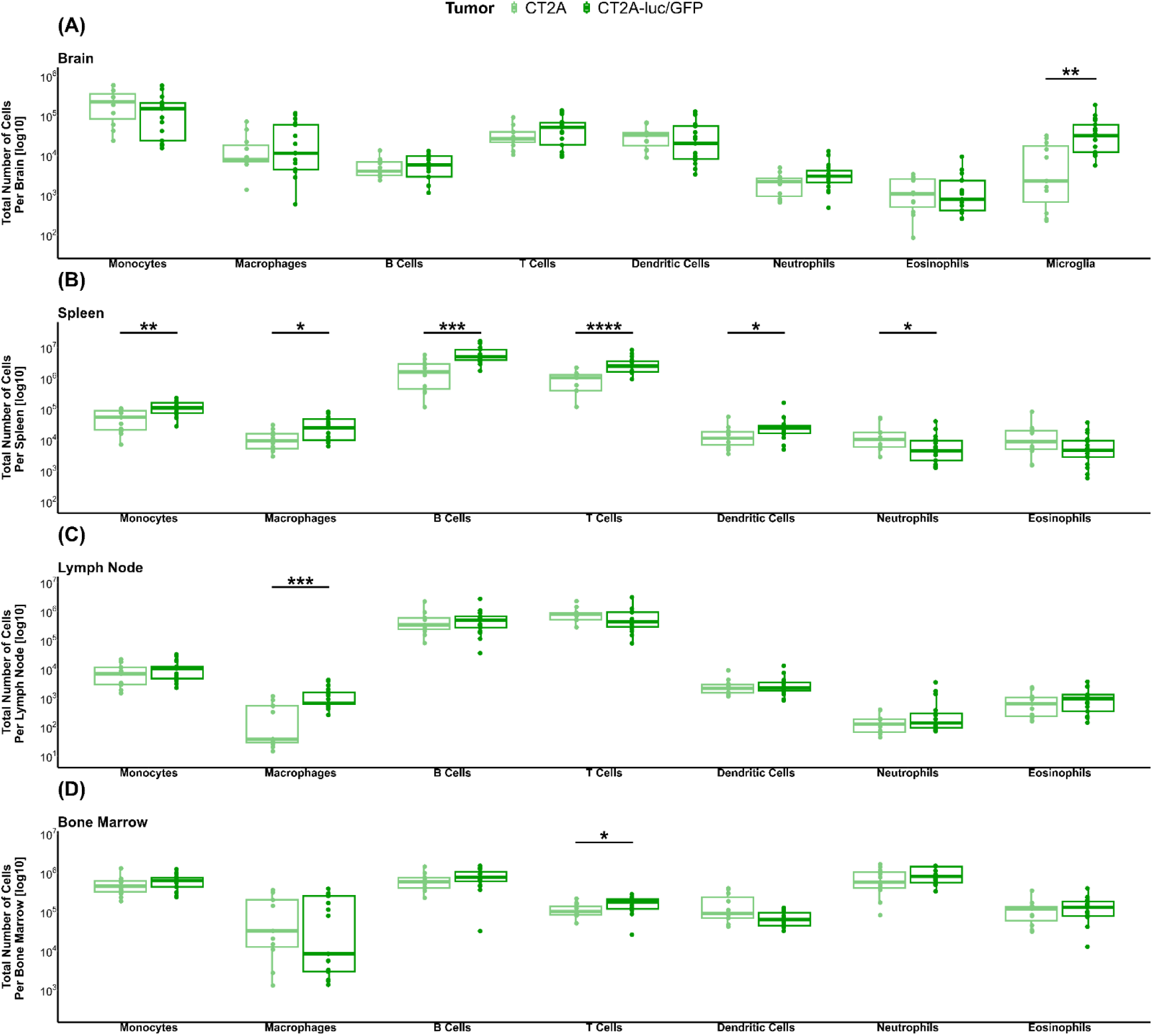
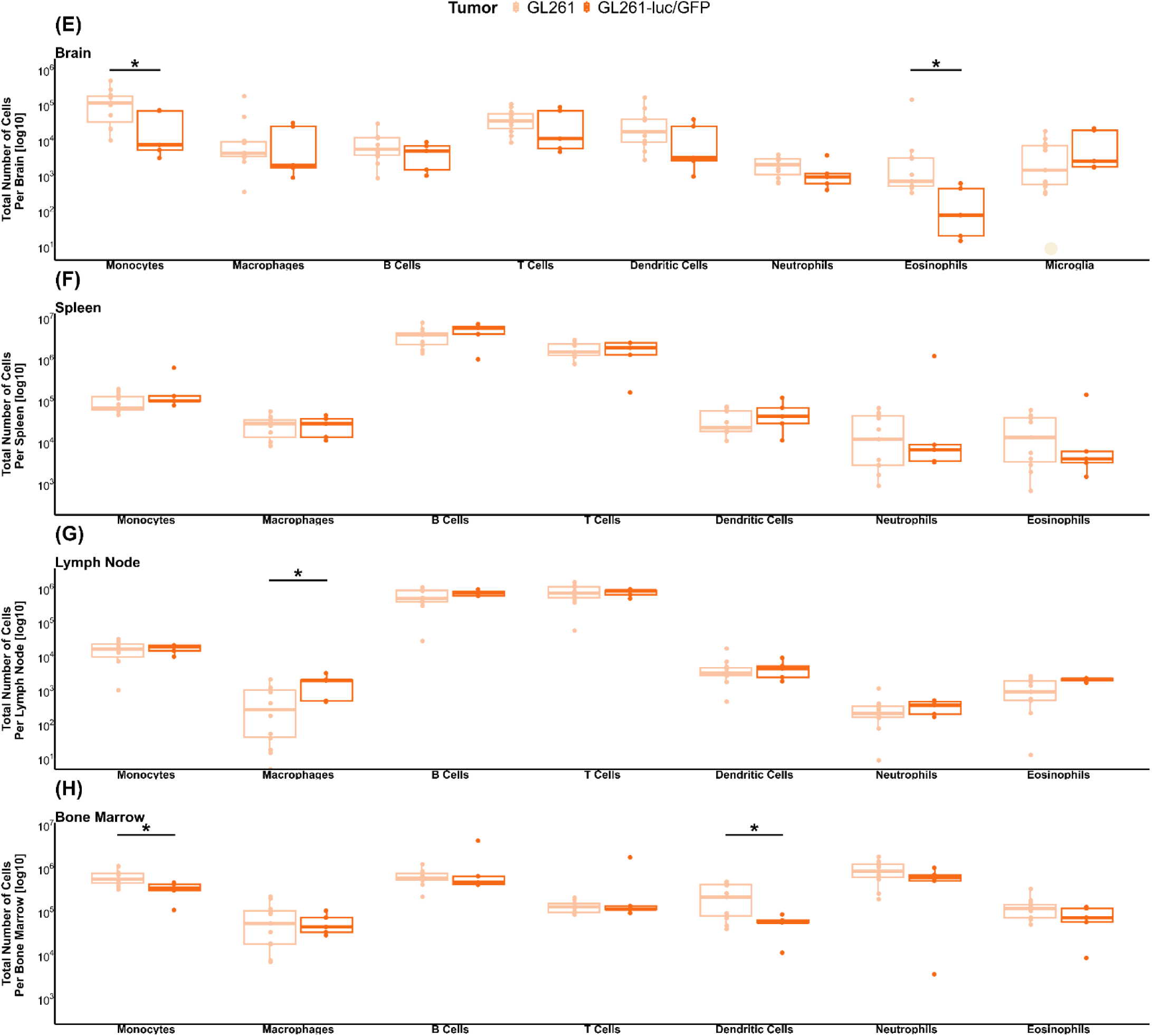
A comparison of immune cells in C57BL/6 mice bearing brain tumors created using CT2A (A-D) or GL261 (E-G) cells \ expressing or not expressing a luciferase/GFP reporter gene cassette. Immune cells were enumerated in the brains (A, E), spleens (B, F) lymph nodes (C, G), and bone marrow (D, H), of mice. Data represent the median, second and third quartiles of n = 11 CT2A tumor-bearing mice, n = 17 CT2A-luc/GFP tumor-bearing mice, n = 11 GL261 tumor-bearing mice, n = 5 GL261-luc/GFP tumor-bearing mice P values *p<0.05, **P<0.01, ***p<0.001, ****p<0.0001.

**Table S2.** Antibodies that were used for staining tissue samples for the identification of immune cells.

| Antibody | Conjugate | Clone | Supplier | Catalogue # |
| --- | --- | --- | --- | --- |
| Viability | FVS 780 |  | BD Biosciences | 565388 |
| CD45 | BUV395 | 30-F11 | BD Biosciences | 564279 |
| FcεR1α | BUV496 | MAR-1 | BD Biosciences | 751763 |
| CD172a | BV750 | P84 | BD Biosciences | 747007 |
| CD11b | BV650 | M1/70 | BD Biosciences | 563402 |
| CD19 | BUV563 | 1D3 | BD Biosciences | 749028 |
| TCR b | BB700 | H57-597 | BD Biosciences | 745846 |
| CD49b | BUV661 | HMα2 | BD Biosciences | 741523 |
| Ly-6G | BV711 | 1A8 | BD Biosciences | 563979 |
| CD45R | V450 | MAR-1 | BD Biosciences | 751763 |
| F4/80 | PE-CF594 | T45-2342 | BD Biosciences | 565613 |
| I-A/I-E (MHC II) | BUV805 | 2G9 | BD Biosciences | 748707 |
| CD11c | BV510 | N418 | BD Biosciences | 744178 |
| Clec9A (CD370) | BB515 | 10B4 | BD Biosciences | 565320 |
| Ly-6C | PE-Cy7 | AL-21 | BD Biosciences | 560593 |
| Siglec-F | APC-R700 | E50-2440 | BD Biosciences | 565183 |
| CD115 | BV605 | T38-320 | BD Biosciences | 743640 |
| PD-L1 (CD274) | BUV615 | MIH5 | BD Biosciences | 752339 |
| CCR7 (CD197) | AF647 | 4B12 | BD Biosciences | 560766 |
| CD86 | BUV737 | PO3 | BD Biosciences | 741757 |
| CD8a | PE | 5H10-1 | BD Biosciences | 567630 |

**Table S3.** Packages utilized in RStudio.

| Package | Version | Reference |
| --- | --- | --- |
| tidyr | 1.3.1 | Wickham H, Vaughan D, Girlich M (2024). *tidyr: Tidy Messy Data*. R package version 1.3.1, https://github.com/tidyverse/tidyr, https://tidyr.tidyverse.org. |
| ggplot2 | 3.5.2 | H. Wickham. *ggplot2: Elegant Graphics for Data Analysis*. Springer-Verlag New York, 2016. |
| dplyr | 1.1.4 | Wickham H, François R, Henry L, Müller K, Vaughan D (2023). *dplyr: A Grammar of Data Manipulation*. R package version 1.1.4, https://github.com/tidyverse/dplyr, https://dplyr.tidyverse.org. |
| ggpubr | 0.6.0 | Kassambara A (2023). *ggpubr: 'ggplot2' Based Publication Ready Plots*. R package version 0.6.0, https://rpkgs.datanovia.com/ggpubr/. |
| ggrepel | 0.9.6 | Slowikowski K (2024). *ggrepel: Automatically Position Non-Overlapping Text Labels with 'ggplot2'*. https://ggrepel.slowkow.com/, https://github.com/slowkow/ggrepel. |
| stringr | 1.5.1 | Wickham H (2023). *stringr: Simple, Consistent Wrappers for Common String Operations*. R package version 1.5.1, https://github.com/tidyverse/stringr, https://stringr.tidyverse.org. |
| rstatix | 0.7.2 | Kassambara A (2023). *rstatix: Pipe-Friendly Framework for Basic Statistical Tests*. R package version 0.7.2, https://rpkgs.datanovia.com/rstatix/. |

## Supplementary Methods

### Patient-Derived Glioblastoma Cell Lines

All GNS cell cultures were passaged when they reached 70–95% confluence by detaching cells using StemPro Accutase (Thermo Fisher) and seeding fresh Matrigel-coated flasks at a 1:3 to 1:5 split ratio. Cultures were grown in a humidified incubator at 37°C with 5% CO_2_.

### C57BL/6 Syngenic Brain Tumour Cell Lines

GD2-GL261-luc/GFP were cultured in T-75 flasks in RPMI medium (Thermo Fisher) supplemented with 10% Foetal Bovine Serum (FBS), 1% Glutamax (Thermo Fisher), 1% Non-Essential Amino Acids (NEAA) (Thermo Fisher), and 1% Penicillin-Streptomycin (PenStrep) (Thermo Fisher). GD2-CT2A-luc/GFP were cultured in T-75 flasks in DMEM medium supplemented with 10% FBS, 1% Glutamax, 1% NEAA, and 1% PenStrep. Both cell lines were passaged when they reached 60–80% confluence by detaching cells using TrypLE (Thermo Fisher) and reseded in fresh flasks at 1:3 to 1:6 ratio.

### Immunofluorescence Staining and Analysis

The GFP and DAPI channels were used to create tumor and non-tumor regions of interest (ROIs). These ROIs were used on the raw unedited CD11b and CD3 channels to measure the mean fluorescence intensity separately for tumor versus non-tumor regions. For the non-tumor-bearing control mice, the fluorescence intensity of expression of CD11b and CD3 were measured in terms of the whole tissue section area.

### Flow Cytometry Staining and Analysis

Cryopreserved dissociated tissue samples were rapidly thawed in a 37°C water bath and transferred to tubes containing 9 mL of warm RPMI. After 10 min at RT, cells were centrifuged and resuspended in FACS buffer (PBS, 0.01%, sodium azide, 0.5% BSA), and Fc receptors were blocked with Mouse Fc Block (BD Biosciences, Cat# 553141) for 10 min at room temperature (RT). Cells were then incubated for 30 min at RT with cocktails containing Brilliant Staining Buffer Plus (BD Biosciences, Cat# 564220), True-Stain Monocyte Blocker (BioLegend, Cat# 421103) and combinations of the antibodies in Table S2. After washing twice in PBS, cells were incubated with a fixable viability stain for 30 min at RT, washed in FACS buffer, resuspended in FACS buffer, and stored at 4° C until acquisition on the same day. Samples were transferred to TruCount tubes (BD Biosciences, Cat# 340334) to enable absolute cell numbers to be determined.

### Immunofluorescence Staining and Analysis

Sections were rehydrated in PBS for 5 minutes at RT and then blocked using 10% normal mouse serum, 10% normal goat serum and 10% normal donkey serum in PBS containing 1% bovine serum albumin (BSA) for 30 minutes at RT.Following washing, sections were incubated with 0.5µg/mL DAPI (BD Pharmingen) in PBS for 5 minutes at RT. Sections were imaged with a Zeiss Axio Scan.Z1 slide-scanner using 20x objective and ZEN 3.7 system software. Analysis of CD11b and CD3 staining was performed using FIJI software (ImageJ, National Institutes of Health) false colour was applied to each channel.

## References.

1 Nguyen, H.-M., Guz-montgomery, K., Lowe, D. B. & Saha, D. Pathogenetic features and current management of glioblastoma. Cancers 13, 1–42, doi:10.3390/cancers13040856 (2021).

2 Stupp, R., et al. Radiotherapy plus Concomitant and Adjuvant Temozolomide for Glioblastoma. New England Journal of Medicine 352, 987–996, doi:10.1056/NEJMoa043330 (2005).

3 Baddam, S. R., Kalagara, S., Kuna, K. & Enaganti, S. Recent advancements and theranostics strategies in glioblastoma therapy. Biomedical Materials 18, 052007, doi:10.1088/1748-605X/acf0ab (2023).

4 Maccari, M., et al. Present and Future of Immunotherapy in Patients With Glioblastoma: Limitations and Opportunities. The Oncologist 29, 289–302, doi:10.1093/oncolo/oyad321 (2023).

5 Gardam, B., Gargett, T., Brown, M. P. & Ebert, L. M. Targeting the dendritic cell-T cell axis to develop effective immunotherapies for glioblastoma. Frontiers in Immunology 14, doi:10.3389/fimmu.2023.1261257 (2023).

6 Gargett, T., et al. GD2-targeting CAR-T cells enhanced by transgenic IL-15 expression are an effective and clinically feasible therapy for glioblastoma. J Immunother Cancer 10, e005187, doi:10.1136/jitc-2022-005187 (2022).

7 Haddad, A. F., et al. Mouse models of glioblastoma for the evaluation of novel therapeutic strategies. Neuro-Oncology Advances 3, doi:10.1093/noajnl/vdab100 (2021).

8 Khalsa, J. K., et al. Immune phenotyping of diverse syngeneic murine brain tumors identifies immunologically distinct types. Nature Communications 11, 3912, doi:10.1038/s41467-020-17704-5 (2020).

9 Himes, B. T., et al. Immunosuppression in Glioblastoma: Current Understanding and Therapeutic Implications. Front Oncol 11, 770561, doi:10.3389/fonc.2021.770561 (2021).

10 Chongsathidkiet, P., et al. Sequestration of T cells in bone marrow in the setting of glioblastoma and other intracranial tumors. Nature Medicine 24, 1459–1468, doi:10.1038/s41591-018-0135-2 (2018).

11 Ayasoufi, K., et al. Brain cancer induces systemic immunosuppression through release of non-steroid soluble mediators. Brain 143, 3629–3652, doi:10.1093/brain/awaa343 (2020).

12 Takahashi, T., et al. Enhanced Antibody Responses in a Novel NOG Transgenic Mouse with Restored Lymph Node Organogenesis. Frontiers in Immunology Volume 8 - 2017, doi:10.3389/fimmu.2017.02017 (2018).

13 Shultz, L. D., et al. Human Lymphoid and Myeloid Cell Development in NOD/LtSz-scid IL2Rγnull Mice Engrafted with Mobilized Human Hemopoietic Stem Cells 12. The Journal of Immunology 174, 6477–6489, doi:10.4049/jimmunol.174.10.6477 (2005).

14 Ito, M., et al. NOD/SCID/γcnull mouse: an excellent recipient mouse model for engraftment of human cells. Blood 100, 3175–3182, 10.1182/blood-2001-12-0207 (2002).

15 Grzelak, C. A., et al. Elimination of fluorescent protein immunogenicity permits modeling of metastasis in immune-competent settings. Cancer Cell 40, 1–2, 10.1016/j.ccell.2021.11.004 (2022).

16 Sanchez, V. E., et al. GL261 luciferase-expressing cells elicit an anti-tumor immune response: an evaluation of murine glioma models. Scientific Reports 10, 11003, doi:10.1038/s41598-020-67411-w (2020).

17 Skelton, D., Satake, N. & Kohn, D. B. The enhanced green fluorescent protein (eGFP) is minimally immunogenic in C57BL/6 mice. Gene Therapy 8, 1813–1814, doi:10.1038/sj.gt.3301586 (2001).

18 Liu, Z., Gu, Y., Shin, A., Zhang, S. & Ginhoux, F. Analysis of Myeloid Cells in Mouse Tissues with Flow Cytometry. STAR Protocols 1, 100029, 10.1016/j.xpro.2020.100029 (2020).

19 Choi, B.-R., Johnson, K. R., Maric, D. & McGavern, D. B. Monocyte-derived IL-6 programs microglia to rebuild damaged brain vasculature. Nature Immunology 24, 1110–1123, doi:10.1038/s41590-023-01521-1 (2023).

20 Spiteri, A. G., Wishart, C. L. & King, N. J. C. Immovable Object Meets Unstoppable Force? Dialogue Between Resident and Peripheral Myeloid Cells in the Inflamed Brain. Frontiers in Immunology 11, doi:10.3389/fimmu.2020.600822 (2020).

21 Spiteri, A. G., et al. High-parameter cytometry unmasks microglial cell spatio-temporal response kinetics in severe neuroinflammatory disease. Journal of Neuroinflammation 18, 166, doi:10.1186/s12974-021-02214-y (2021).

22 Shultz, L. D., et al. Multiple defects in innate and adaptive immunologic function in NOD/LtSz-scid mice. The Journal of immunology (1950) 154, 180–191, doi:10.4049/jimmunol.154.1.180 (1995).

23 Pollard, S. M., et al. Glioma stem cell lines expanded in adherent culture have tumor-specific phenotypes and are suitable for chemical and genetic screens. Cell Stem Cell 4, 568–580, doi:10.1016/j.stem.2009.03.014 (2009).

24 Dudek, A. M., Martin, S., Garg, A. D. & Agostinis, P. Immature, Semi-Mature, and Fully Mature Dendritic Cells: Toward a DC-Cancer Cells Interface That Augments Anticancer Immunity. Front Immunol 4, 438, doi:10.3389/fimmu.2013.00438 (2013).

25 Mukherjee, R., et al. Non-Classical monocytes display inflammatory features: Validation in Sepsis and Systemic Lupus Erythematous. Scientific Reports 5, 13886, doi:10.1038/srep13886 (2015).

26 Murty, S., et al. Intravital imaging reveals synergistic effect of CAR T-cells and radiation therapy in a preclinical immunocompetent glioblastoma model. Oncoimmunology 9, 1757360, doi:10.1080/2162402x.2020.1757360 (2020).

27 Weiss, T., Weller, M., Guckenberger, M., Sentman, C. L. & Roth, P. NKG2D-Based CAR T Cells and Radiotherapy Exert Synergistic Efficacy in Glioblastoma. Cancer Res 78, 1031–1043, doi:10.1158/0008-5472.Can-17-1788 (2018).

28 Donlon, N. E., Power, R., Hayes, C., Reynolds, J. V. & Lysaght, J. Radiotherapy, immunotherapy, and the tumour microenvironment: Turning an immunosuppressive milieu into a therapeutic opportunity. Cancer Letters 502, 84–96, 10.1016/j.canlet.2020.12.045 (2021).

29 Tennant, M. D., New, C., Ferreira, L. M. R. & O’Neil, R. T. Efficient T cell adoptive transfer in lymphoreplete hosts mediated by transient activation of Stat5 signaling. Molecular Therapy 31, 2591–2599, doi:10.1016/j.ymthe.2023.07.015 (2023).

30 Johnson, C. B., et al. Enhanced Lymphodepletion Is Insufficient to Replace Exogenous IL2 or IL15 Therapy in Augmenting the Efficacy of Adoptively Transferred Effector CD8+ T Cells. Cancer Research 78, 3067–3074, doi:10.1158/0008-5472.Can-17-2153 (2018).

31 Ko, Y., Jeong, Y. H. & Lee, J. A. Effects of Low- or Moderate-dose Whole Body-X-ray Radiation on the Immune System of C57BL/6 Mice. Clinical pediatric hematology-oncology 25, 50–55, doi:10.15264/cpho.2018.25.1.50 (2018).

32 Bogdándi, E. N., et al. Effects of Low-Dose Radiation on the Immune System of Mice after Total-Body Irradiation. Radiation research 174, 480–489, doi:10.1667/RR2160.1 (2010).

33 Liu, Y., Zhou, F., Ali, H., Lathia, J. D. & Chen, P. Immunotherapy for glioblastoma: current state, challenges, and future perspectives. Cellular & Molecular Immunology 21, 1354–1375, doi:10.1038/s41423-024-01226-x (2024).

34 Salvato, I. & Marchini, A. Immunotherapeutic Strategies for the Treatment of Glioblastoma: Current Challenges and Future Perspectives. Cancers 16, 1276 (2024).

35 Omuro, A., et al. Nivolumab with or without ipilimumab in patients with recurrent glioblastoma: results from exploratory phase I cohorts of CheckMate 143. Neuro-oncology 20, 674–686 (2018).

36 Omuro, A., et al. Nivolumab plus radiotherapy with or without temozolomide in newly diagnosed glioblastoma: Results from exploratory phase I cohorts of CheckMate 143. Neuro-oncology advances 4, vdac025-vdac025, doi:10.1093/noajnl/vdac025 (2022).

37 Agosti, E., et al. Glioblastoma immunotherapy: A systematic review of the present strategies and prospects for advancements. International Journal of Molecular Sciences 24, 15037 (2023).

38 Yeo, E. C. F., Brown, M. P., Gargett, T. & Ebert, L. M. The Role of Cytokines and Chemokines in Shaping the Immune Microenvironment of Glioblastoma: Implications for Immunotherapy. *Cells (Basel*, Switzerland) 10, 607, doi:10.3390/cells10030607 (2021).

39 Sharma, P., Aaroe, A., Liang, J. & Puduvalli, V. K. Tumor microenvironment in glioblastoma: Current and emerging concepts. Neurooncol Adv 5, vdad009, doi:10.1093/noajnl/vdad009 (2023).

40 Gieryng, A., Pszczolkowska, D., Walentynowicz, K. A., Rajan, W. D. & Kaminska, B. Immune microenvironment of gliomas. Laboratory Investigation 97, 498–518, doi:10.1038/labinvest.2017.19 (2017).

41 Gousias, K., von Ruecker, A., Voulgari, P. & Simon, M. Phenotypical analysis, relation to malignancy and prognostic relevance of ICOS + T regulatory and dendritic cells in patients with gliomas. Journal of neuroimmunology 264, 84–90, doi:10.1016/j.jneuroim.2013.09.001 (2013).

42 Löhr, M., et al. High-grade glioma associated immunosuppression does not prevent immune responses induced by therapeutic vaccines in combination with T reg depletion. Cancer immunology, immunotherapy (2018).

43 Sheu, T.-T. & Chiang, B.-L. Lymphopenia, Lymphopenia-Induced Proliferation, and Autoimmunity. International journal of molecular sciences 22, 4152, doi:10.3390/ijms22084152 (2021).

44 Grossman, S. A., et al. Immunosuppression in Patients with High-Grade Gliomas Treated with Radiation and Temozolomide. Clinical Cancer Research 17, 5473–5480, doi:10.1158/1078-0432.Ccr-11-0774 (2011).

45 Ghosh, S., et al. Radiation-induced circulating myeloid-derived suppressor cells induce systemic lymphopenia after chemoradiotherapy in patients with glioblastoma. Science translational medicine 15, eabn6758, doi:10.1126/scitranslmed.abn6758 (2023).

46 Maddison, K., et al. Low tumour-infiltrating lymphocyte density in primary and recurrent glioblastoma. Oncotarget 12, 2177–2187, doi:10.18632/oncotarget.28069 (2021).

47 Gargett, T. & Brown, M. P. Different cytokine and stimulation conditions influence the expansion and immune phenotype of third-generation chimeric antigen receptor T cells specific for tumor antigen GD2. *Cytotherapy (Oxford*, England) 17, 487–495, doi:10.1016/j.jcyt.2014.12.002 (2015).

48 Gargett, T., et al. Safety and biological outcomes following a phase 1 trial of GD2-specific CAR-T cells in patients with GD2-positive metastatic melanoma and other solid cancers. J Immunother Cancer 12, e008659, doi:10.1136/jitc-2023-008659 (2024).

49 Tirapelle, M. C., Schmidt, D. D., Ebrahimabadi, S., Covas, D. & Picanço-Castro, V. GD2-TARGETING CAR-NK CELLS ENHANCED BY TRANSGENIC GITRL EXPRESSION ARE AN EFFECTIVE FOR GLIOBLASTOMA AND MELANOMA. *Cytotherapy (Oxford*, England) 26, S199–S199, doi:10.1016/j.jcyt.2024.03.398 (2024).

50 Chiavelli, C., et al. Autologous anti-GD2 CAR T cells efficiently target primary human glioblastoma. NPJ PRECISION ONCOLOGY 8, 26–26, doi:10.1038/s41698-024-00506-z (2024).

51 Richman, S. A., et al. High-Affinity GD2-Specific CAR T Cells Induce Fatal Encephalitis in a Preclinical Neuroblastoma Model. Cancer Immunology Research 6, 36–46, doi:10.1158/2326-6066.Cir-17-0211 (2018).

52 Tang, L., Pan, S., Wei, X., Xu, X. & Wei, Q. Arming CAR-T cells with cytokines and more: Innovations in the fourth-generation CAR-T development. Molecular Therapy 31, 3146–3162, doi:10.1016/j.ymthe.2023.09.021 (2023).

53 Ageenko, A., Vasileva, N., Richter, V. & Kuligina, E. Combination of Oncolytic Virotherapy with Different Antitumor Approaches against Glioblastoma. International Journal of Molecular Sciences 25, 2042 (2024).

54 Segura-Collar, B., et al. Advanced immunotherapies for glioblastoma: tumor neoantigen vaccines in combination with immunomodulators. Acta Neuropathologica Communications 11, 79, doi:10.1186/s40478-023-01569-y (2023).

55 Houweling, M., et al. Screening of predicted synergistic multi-target therapies in glioblastoma identifies new treatment strategies. Neuro-Oncology Advances 5, doi:10.1093/noajnl/vdad073 (2023).

56 Ebert, L. M., et al. Endothelial, pericyte and tumor cell expression in glioblastoma identifies fibroblast activation protein (FAP) as an excellent target for immunotherapy. Clinical & Translational Immunology 9, e1191, 10.1002/cti2.1191 (2020).

57 Thomas, S., Straathof, K., Himoudi, N., Anderson, J. & Pule, M. An Optimized GD2-Targeting Retroviral Cassette for More Potent and Safer Cellular Therapy of Neuroblastoma and Other Cancers. PLoS One 11, e0152196, doi:10.1371/journal.pone.0152196 (2016).

58 Ullman-Culleré, M. H. & Foltz, C. J. Body condition scoring: a rapid and accurate method for assessing health status in mice. Lab Anim Sci 49, 319–323 (1999).

